# Rapid Characterization of AAV gene therapy vectors by Mass Photometry

**DOI:** 10.1101/2021.02.18.431916

**Authors:** Di Wu, Philsang Hwang, Tiansen Li, Grzegorz Piszczek

**Affiliations:** Biophysics Core Facility, National Heart, Lung, and Blood Institute, 50 South Drive, Bethesda, MD 20892-8012, USA; Ocular Gene Therapy Core Facility, National Eye Institute, 6 Center Drive, Bethesda, Maryland 20892, USA

## Abstract

Recombinant adeno-associated viruses (rAAV) are extensively used as gene delivery vectors in clinical studies, and several rAAV based treatments have already been approved. Significant progress has been made in rAAV manufacturing, and large-scale vector production and purification methods have been developed. However, a better and more precise capsid characterization techniques are still needed to guarantee the purity and safety of the rAAV preparations. A recently developed single-molecule technique, mass photometry (MP), measures mass distributions of biomolecules with high resolution and sensitivity. Here we explore applications of MP for the characterization of capsid fractions. We demonstrate that MP is able to resolve and quantify not only empty and full-genome containing capsid populations, but also identify the partially packaged capsid impurities. MP data accurately measures full and empty capsid ratios, and can be used to estimate the size of the encapsidated genome. MP distributions provide information on sample heterogeneity and on the presence of aggregates. Current analytical techniques used to characterize rAAV preparations are susceptible to background signals, have limited accuracy, or are time-consuming and require a large amount of material. MP can analyze sub-picomole quantities of sample, and data can be obtained and analyzed within minutes. This method provides a simple, robust, and effective tool to monitor physical attributes of rAAV vectors.

## Introduction

Adeno-associated viruses (AAVs) are small viruses that infect humans and non-human primates. Several features make recombinant AAVs (rAAVs) attractive as a tool for gene therapy applications. AAVs lack pathogenicity, their replication depends on coinfection with a helper virus, and they can carry their genetic material to both dividing and quiescent cells [1, 2]. The first rAAV based therapy was approved by the US Food and Drug Administration in 2012, and five viral vector treatments are currently available [3]. There are also over 200 ongoing clinical trials for various rAAV based gene therapies [4]. The increasing demand for rAAVs for both laboratory [5, 6] and clinical applications accelerated the development of new techniques for rAAV production and characterization. Considerable progress has been made in the large-scale production and purification of the rAAV vectors, but there is still a need for new analytical methods to characterize and quantify rAAV preparations. Complex biotherapeutic products are inherently heterogenous and difficult to characterize, but gene therapy vectors pose additional unique challenges. One of them is the high complexity of viral vectors, consisting of the protein capsid assembly and the DNA genome. Particularly challenging is the presence of impurities that closely resemble the desired product, namely empty capsids, and capsids carrying nucleic acid impurities. An ideal analytical method should be able to not only overcome those challenges, but also obtain data quickly and with minimal sample consumption, while delivering robust results that can be validated and standardized to meet biotherapeutic production regulatory requirements.

The wild-type AAV contains a single-stranded DNA (ssDNA) approximately 4.7 kb long, encoding the replication (*rep*) and encapsidation (*cap*) genes, flanked by inverted terminal repeats (ITRs) [7, 8]. The AAV genome is contained within an approximately 26 nm diameter capsid, consisting of 60 protein molecules [9, 10]. rAAVs have the same capsid structure, but the (*rep*) and (*cap*) genes are removed, and the gene of interest is inserted between the ITRs. The rAAVs are typically produced by co-transfecting the host cells with three plasmids: a plasmid containing the gene of interest and two helper plasmids that eliminate the need for the adenoviral coinfection [11-13]. During the virion assembly, the DNA genome is inserted into the preassembled capsid. Consequently, besides the properly assembled virions, the final product can also contain non-genome-containing particles (empty capsids) and vectors containing incomplete portions of the genome or nongenomic nucleic acid contaminants [14, 15].

Depending on the particular vector production method and the product type, empty capsids concentration can be over 20-times higher than the genome containing vectors [16]. The effect of empty capsid impurities on the gene therapy clinical outcomes is not fully understood [17, 18], but there is little doubt that the development of reliable methods for precise characterization of all viral product impurities is critically important.

Over the years, several techniques have been applied for rAAV product characterization. Technologies previously validated for other biologics can be used for the analysis of some rAAV impurities, such as the residual host cell protein contamination, but often are not able to detect and quantify the empty and partially filled capsid populations. In particular, the diffusion-based methods, such as size exclusion chromatography or dynamic light scattering, are not suitable for vector packing efficiency determination since both the empty capsids and the full vectors have identical geometric size. Typically, the ratio of empty to full virions is assessed by negative-stain transmission electron microscopy (TEM), optical density measurements, or by the combination of quantitative PCR (qPCR) and ELISA [19-21]. Despite their routine use, all three methods tend to be imprecise, and none of the three are able to reliably quantify the partially filled virions. In optical density and TEM measurements, the background contributes to the signal, and the negative stain can affect the particles under TEM analysis. The combination of qPCR and ELISA compounds errors of the two methods, and is also dependent on the antibodies and amplicons applied in the ELISA and qPCR analyses, respectively. Improved implementations of those methods have been developed: for example, the combination of UV measurements with multi-angle light scattering and size exclusion chromatography (SEC-UV-MALS) reduces background contributions [22], and Cryo-electron microscopy (CryoEM) eliminates negative staining [23]. Neither SEC-UV-MALS nor CryoEM, however, are sensitive to the partially filled virion populations. Ion-exchange chromatography has been successfully applied to separate empty and full vectors [16, 24] and its scalability makes it particularly useful for the empty vector removal step in the rAAV production process. This method separates particles using differences in the isoelectric points of viral fractions containing different amounts of the strongly negatively-charged DNA in their capsids.

However, despite its effectiveness for the empty and full-vector separation, ion-exchange chromatography does not usually have a high enough resolution to analyze the partially filled populations. The two currently available methods effective in the detection and quantification of all viral fractions, including the empty, partially filled, and full capsids, are analytical ultracentrifugation (AUC), and charge detection mass spectrometry (CDMS) [25, 26]. AUC provides a high degree of resolution of viral species based on differences in their buoyant masses. Additionally, the optical density and refractive index detection systems of analytical ultracentrifuge allow for the estimation of the protein to DNA ratio for all viral species. Neither AUC nor CDMS are widely used due to the large amount of sample and long analysis time required by AUC and the lack of commercial instrument availability of CDMS.

Here, we apply mass photometry (MP), a recently developed single-molecule technique based on interferometric scattering microscopy (iSCAT) [27, 28] to the analysis of the rAAV preparations. MP detects unmodified molecules in solution when they land on the surface of a microscope coverslip. The local contrast changes in the microscope image that originate from the light scattered by individual molecules are measured by the camera (supplemental Fig. 1). This signal is proportional to the molecular masses of the scattering particles. MP analysis yields the particle mass distributions histogram, comprised of several thousand data points representing individually detected molecules. The native MP distributions are obtained in contrast units, but since MP contrast is proportional to molecular mass, contrast distributions can easily be converted to mass distributions. Contrast-to-mass (CTM) calibration values for this conversion are obtained from measurements of the appropriate molecular mass standards. To date, MP has been used for the analysis of proteins and their complexes [29-32], as well as larger structures, including protein cages and viral capsids [33-35]. MP can detect molecules in the molecular mass range from approximately 40 kDa to 10 MDa. A typical measurement requires only 10 μL of sample volume at a concentration of approximately 20 nM, and data acquisition and analysis can be completed in less than 5 minutes. This combination of sensitivity and speed makes MP a potentially very attractive tool for the assessment of purity, composition and homogeneity of the rAAV preparations.

To test MP performance, and to assess the quality of the mass photometry rAAV data, we conducted a detailed study evaluating a variety of rAAV samples and validated the MP results with sedimentation velocity AUC (SV-AUC) analysis. Highly purified, genome-containing rAAVs and empty capsid preparations were used to test MP resolution, repeatability, and the ability to quantify the sample fractional content. Vectors with different genome sizes were used to obtain the MP contrast-to-genome-size calibration. Finally, partially purified, commercial rAAV preparations were tested to assess the information content of the MP distributions for more complex, lower purity samples. We demonstrate that MP is a robust, stereotype-independent rAAV analysis method. MP data can be obtained and analyzed very quickly, using a sub-picomole amount of sample. MP provides information on sample heterogeneity and on the relative fractional content of rAAV populations. Additionally, MP identifies small amounts of the partially filled capsids—an impurity often left undetected by other analytical methods. Contrast values provided by the MP distributions can also be used to estimate the genome size of the analyzed virions. More work will have to be done to fully validate and standardize MP for clinical product analysis, but data presented here show that MP has a potential to become one of the standard tools for the rAAV quality assessment.

## Materials and Methods

### Production and purification of rAAV vectors

The rAAV8 vectors were obtained by the calcium phosphate transfection method as described previously [36]. Briefly, the HEK293 cells (ATCC CRL-1,573; ATCC, Manassas, VA) were seeded in vented cap roller bottles (850 cm^2^; Corning, New York, NY) and transfected with 150 µg each of the three plasmids: ITR-containing vector plasmid, Adenoviral helper plasmid for AAV replication, and the AAV replication/capsid proteins plasmid. The vector-containing cells were harvested after 48 hours and disrupted by microfluidization (model HC 2000; Microfluidics Corporation, Newton, MA). After a series of centrifugation steps to eliminate the tissue debris, the supernatant was digested with Benzonase (100 U/ml). Vector particles were congregated with 8% polyethylene glycol 8000 and dissolved in HEPES buffer with RNase A (50 mM HEPES, 150 mM NaCl, 20 mM EDTA, 1% Sarkosyl, pH 8.0, 10 µg/ml RNase A). A cesium chloride step gradient followed by ultracentrifugation was performed to purify the vector-containing dissolvent. Bands containing vector fractions were identified by visual inspection, drawn using 18-gauge hypodermic needle, diafiltered against Tris-buffered saline (10 mM Tris-Cl, 180 mM NaCl, pH 7.4) with 0.001% Pluronic F68, and stored at -80°C. The vector genome titers were determined by qPCR using primers against the promoters or by the SDS-PAGE with Silver staining. The scAAV5-hSyn-GFP and scAAV5-CMV-GFP vectors (Fig. 5E-H) were obtained from Virovek (Hayward, CA) and used without further purification.

### Concentration determination of the rAAV stocks

To prepare mixtures with precisely defined fractions of the empty and full capsids (rAAV8 and rAAV8_CMV_EGFP, respectively), the UV absorbance of the purified vector stocks was acquired using a NonoDrop-2000 spectrophotometer (Thermofisher, Wattham, MA). The extinction coefficient of AAV capsids is 6.61 ×10^6^ M^-1^cm^-1^ at 280 nm (*ε*_cap280_), and 3.72 ×10^6^ M^-1^cm^-1^ at 260 nm (*ε*_cap260_), respectively. Similarly, the extinction coefficients of DNA molecules is 11.1 g^-1^cm^-1^ at 280 nm (*ε*_DNA280_), and 20 g^-1^cm^-1^ at 260 nm (*ε*_DNA260_), respectively [21]. The observed absorbance values are given by:

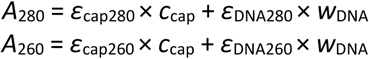

where *c*_cap_ is the capsid molar concentration and *w*_DNA_ is the w/v concentration of the DNA. The average values of 3.74 MDa for the AAV capsid mass, and 650 Da for the DNA base pair MW, respectively, were used for the vector particle concentration calculations.

### MP measurements

The MP experiments were performed at room temperature using the OneMP instrument (Refeyn, UK). The 24×50 mm microscope coverslips (Fisher Scientific, Waltham, MA) were prepared by cleaning with MilliQ water and isopropanol, and drying under a stream of clean nitrogen, as described previously [37]. A piece of clean, precut 2×2-well culturewell gasket (GBL103250, Sigma, MO) was attached to the coverslip. The rAAV stocks were diluted in PBS to a concentration of about 10^11^ virus particles per milliliter. Ten microliters of the filtered PBS buffer were loaded into a well of the culturewell gasket, and, after MP focusing, a 10 µL of rAAV sample was added into the same well. Immediately after the solution was mixed by pipetting, a 2-minute video was recorded using the AcquireMP (Refeyn, UK) software.

### MP data analysis

The MP video files were processed using the DiscoverMP software (Refeyn, UK). The MP detection range is approximately 40 kDa to 10 MDa, and the rAAV samples analyzed here fall into the 3 to 5 MDa range. During data processing, the “Filter-1” value was set to 5 to focus the analysis on the high-MW particle range. The Filter-1 sets the threshold for a minimum step change in the signal value used to detect the molecule landing events. Higher numerical values of Filter-1 effectively increase the low-MW detection cutoff of MP. MP contrast distributions were plotted as Kernel Density Estimates (KDE) or histograms. For all plots, the KDE contrast bandwidth and the histogram bin size were both set to 0.003 contrast units. To obtain information on the contrast distribution species, the MP histograms were fit with Gaussian peaks. For each fitted species, the best fit Gaussian peak position and area represent their average contrast value and their number fraction, respectively. To estimate the molecular mass of the empty capsid sample, the MP contrast distribution was converted to a mass distribution by applying the CTM calibration obtained using an unstained protein ladder sample (LC0725, Thermofisher, Wattham, MA)

### AUC analysis

The AUC experiments were performed in the Proteome Lab XLI (Beckman Coulter, Indianapolis, IN), using the 12 mm pathlength, double sector cells with charcoal filled Epon centerpieces and sapphire windows. The rAAV sample concentrations were adjusted to obtain approximately 0.2 OD_260 nm_ absorbance signals at a 1.2 cm pathlength. Sample and reference channels were each loaded with 400 μL of the rAAV solution and the PBS buffer, respectively. Loaded cells were placed in the 4-hole analytical rotor and allowed to equilibrate thermally in the centrifuge at 20°C, under vacuum, for 60 minutes. After thermal equilibrium was reached, the rotor was accelerated to 10,000 rpm and the Rayleigh interference and 260 nm absorbance scans in the intensity mode were started immediately. Radial scans were collected continuously until the sedimenting boundary cleared the detection window. Data analysis was performed in the SEDFIT software (National Institutes of Health) using the *c*(*s*) distribution model [38].

## Results

### MP analysis of empty and genome-containing rAAV vectors

To test the applicability of MP to rAAV characterization, we started with the analysis of the purified rAAV fractions. Concentrations of the genome-containing viral vectors (rAAV_gc) and the empty viral particle (rAAV_ec) stock solutions were determined by the A_260_/A_280_ absorption (see Methods section). For the MP analysis, all final sample concentrations were adjusted to approximately 1 × 10^11^ particles/mL. The MP contrast distributions of both the rAAV_ec and rAAV_gc show single, distinct distribution peaks (Fig. 1A). The rAAV_ec peak position is shifted to smaller contrast values in respect to the rAAV_gc peak, reflecting a smaller total mass of the empty viral particles. Additionally, the rAAV_ec peak is relatively narrow, whereas the increased width of the rAAV_gc peak indicates higher heterogeneity of the rAAV_gc preparation. Slight fluctuations of the distribution baselines outside of the peak areas may indicate the presence of a small amount of particle fragments, impurities, or aggregates. The molecular mass of empty capsids calculated from the average contrast of the rAAV_ec peak using the CTM calibration based on the protein mass standards is 3.72 ±0.15 MDa which is in very good agreement with the predicted molecular mass of 3.74 MDa [21]. Replicates of the MP measurements show only small fluctuations of the rAAV_ec peak position, indicating good repeatability of the MP analysis (Fig. 2).

**Figure 1.**
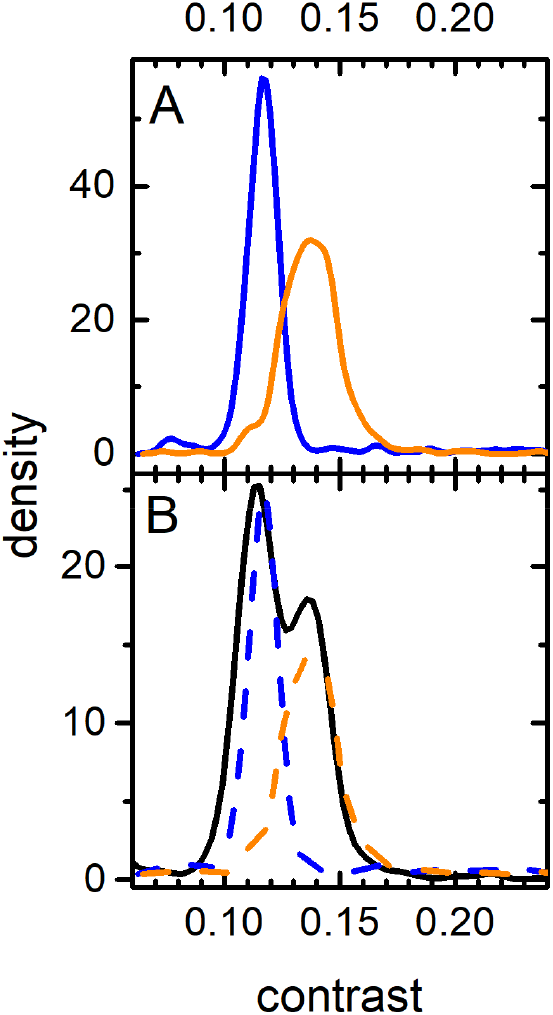
KDE mass distributions of the purified rAAV preparations. (A) Mass distributions of the empty capsids (rAAV8, in blue) and genome containing vectors (AAV8-CMV-EGFP, in orange). (B) Mass distribution of the empty and genome containing rAAVs mixture. Dashed lines represent the individually measured components shown in panel A, rescaled to account for sample dilution.

**Figure 2.**
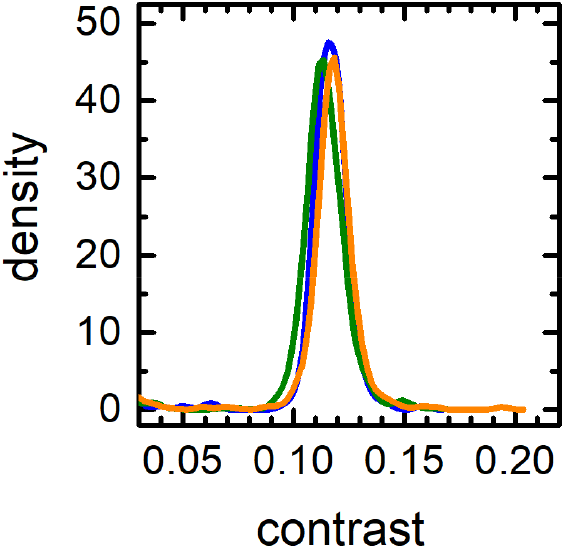
KDE plot of the triplicate MP measurements of the empty rAAV capsid sample shown in Figure 1A.

To evaluate the MP capability to discriminate empty and full capsid populations in a complex sample, equal volumes of the rAAV_ec and rAAV_gc stocks were mixed before dilution for the MP measurement. The MP contrast distribution obtained for the mixed sample shows two distinct peaks representing two rAAV components (Fig. 1B). The position of each peak corresponds to the respective contrast values obtained from the analysis of rAAV_ec and rAAV_gc samples (Fig. 1B, dotted lines). These results show that MP measurements can quickly identify, and assess the quality of, purified AAV fractions. Furthermore, empty and genome containing rAAV particle populations can be effectively discerned from the MP distributions.

### Quantification of empty and genome-containing vector fractions

The number of molecules detected for each species in the MP distribution is given by the area of the corresponding distribution peak. Moreover, the protein population ratios obtained from MP distributions faithfully represent the ratios of their solution concentrations [22]. We applied the same strategy to characterize relative populations in the rAAV mixtures. The empty and full vector fractions in a rAAV sample (Fig. 1B) were quantified by fitting the MP distribution histogram with two Gaussian functions (Fig. 3A). The rAAV_ec to rAAV_gc ratio obtained from the Gaussian peak areas is 1.30, in good agreement with the expected ratio of 1.26, calculated from the concentrations of the rAAV stocks. To further verify this finding, we measured the MP distributions of several mixtures of empty and full rAAV samples at different species mixing ratios (Fig. 3B). Four MP measurements were taken for each sample, and the vertical error bars represent the standard deviation of the measurements. The abscissa values were calculated from the known rAAV_ec and rAAV_gc stock concentrations as determined by the UV absorbance. Values obtained from the MP analysis are in excellent agreement with the expected empty to full fraction ratios (slope = 1.01 ±0.02, R^2^ = 0.997). This indicates that the MP not only detects and identifies the empty and full capsid species in a rAAV sample but can also be used as a powerful tool to quickly determine their relative concentrations.

**Figure 3.**
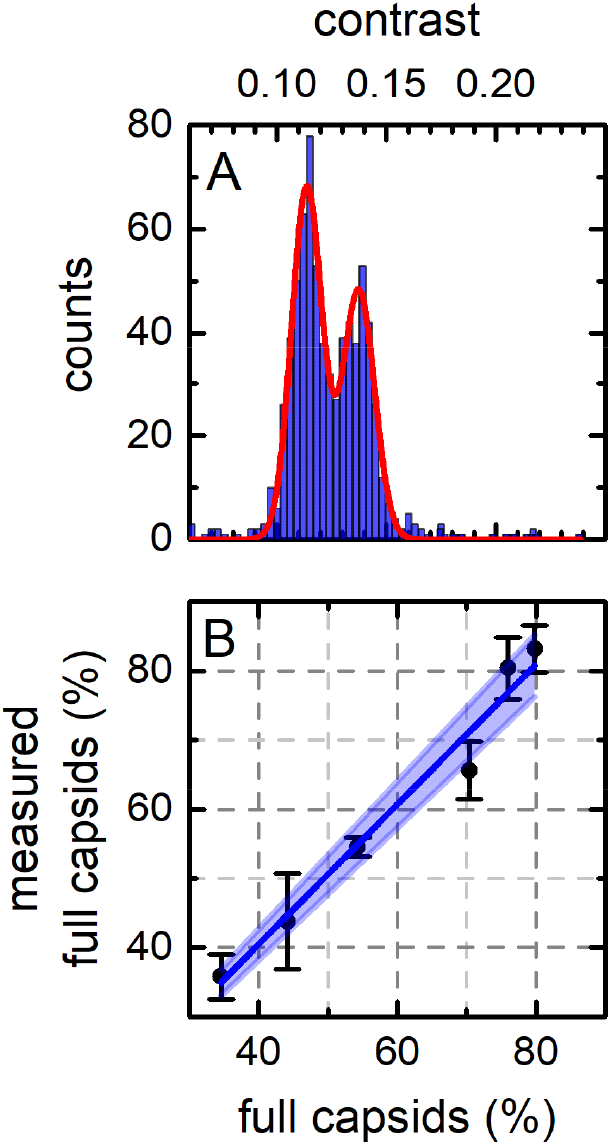
MP determination of the full to empty capsid ratios. (A) MP mass distribution histogram (blue) and the Gaussian fit (red) of the full and empty capsid mixture shown in Fig. 1B. (B) Full capsid fraction determined from the MP measurements plotted against the expected fraction obtained from stocks mixing ratios. Vertical error bars represent the standard deviations of four MP measurements. The dark blue line represents a linear fit (R^2^ = 0.997), and the shaded area indicates the 95%CI band.

### Estimation of the rAAV genome size

The MP signal originates from the interference of the light scattered by molecules binding to the coverslip surface and the light reflected at the glass-water interface. In complex viral particles, both the protein and nucleic acid components contribute to the light scattering signal [28, 39]. Since, for the rAAV virions containing genomes of different sizes, the capsid component signal contributions are identical, the differences in the observed MP contrast values should reflect the different sizes of the encapsidated genomes. Here, we tested several rAAV preparation containing different sizes of both single- and double-stranded DNA genomes. Figure 4 shows the experimental contrast values of five rAAVs containing single-stranded DNA (Fig. 4A) and two rAAV constructs with double-stranded, self-complementary DNA genomes (Fig. 4B), plotted as a function of their genome size. Each measurement was repeated seven times, and the vertical error bars represent standard deviations of the average contrast values. Values obtained for empty vectors (zero genome size) were included on both plots. Both data sets show a linear relationship between the MP contrast and the DNA length with R^2^ of 0.957 and 0.994 for the ssDNA and dsDNA, respectively. The fit parameters define the contrast-to-nucleotide ratios that can be used as calibration values to determine the rAAV genome size from MP measurements. To compare the contrast-to-nt value obtained for the ssDNA to the contrast-to-bp value of dsDNA, the former has to be multiplied by 2, resulting in the value of 0.0199 ±0.0018/kb. This value is higher than the 0.0152 ±0.0008/kb obtained for the dsDNA. This observation is consistent with the previous findings [29], where differences were observed for the contrast-to-nt relationships obtained for the isolated ss- and dsDNA. Since slope differences observed here for the ss- and dsDNA containing rAAVs are small, all experimental data points were combined and plotted as a function of the DNA length (Fig. 4C). The linear plot shows a direct proportionality of the experimental MP contrast values and the rAAV genome sizes for the whole range of tested DNA lengths. No outliers were detected, but the R^2^ value of 0.908 for the combined data plot is lower than that obtained from the individual analysis of the single- and double-stranded DNA data sets. The linear relationships observed here expand the applicability of MP analysis to the measurements of the sizes of genomes packaged into rAAV vectors.

**Figure 4.**
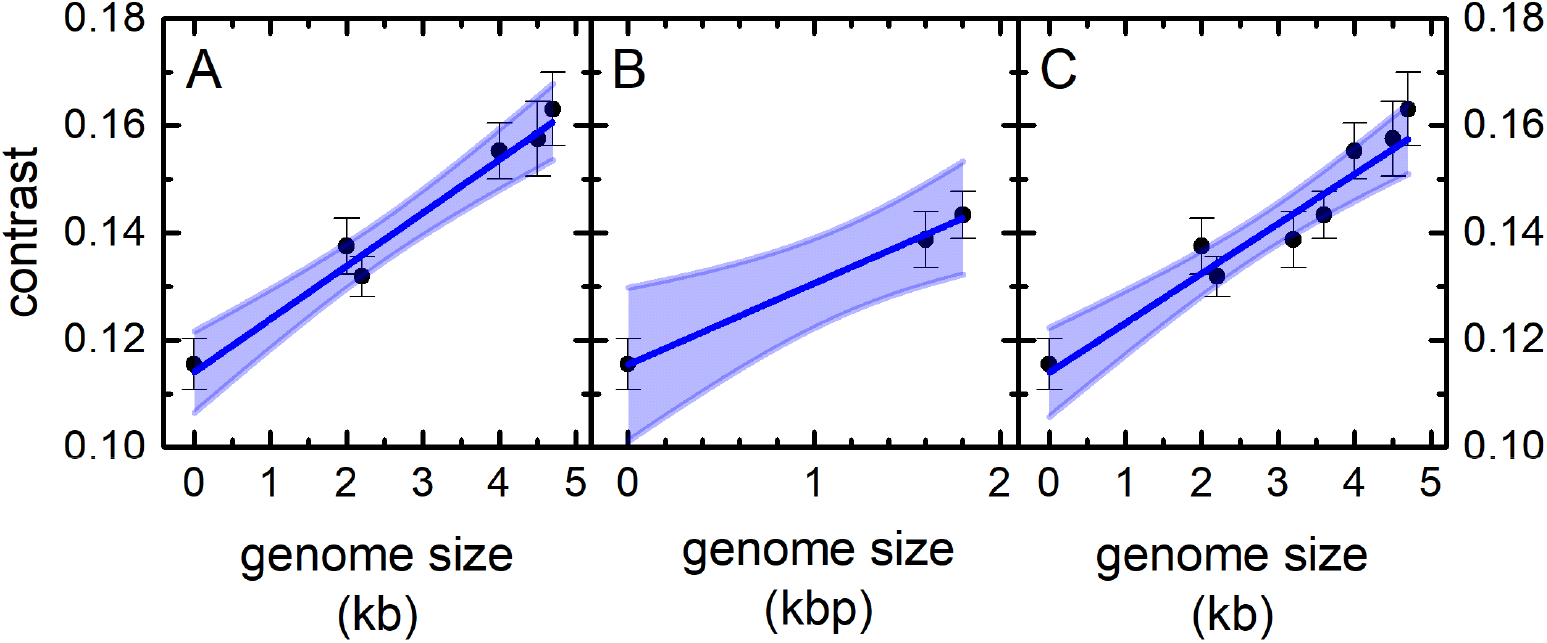
MP distributions peak positions plotted against the sample genome size. (A) ssDNA and (B) dsDNA genome vectors. (C) Combined plot of the ssDNA and dsDNA vector data. Vertical error bars represent the standard deviations of seven MP measurements. Dark blue lines represent linear fits, with R^2^ of 0.957, 0.994, 0.908 for lines in panel (A), (B), and (C), respectively. Shaded areas indicate the 95% CI bands.

**Figure 5.**
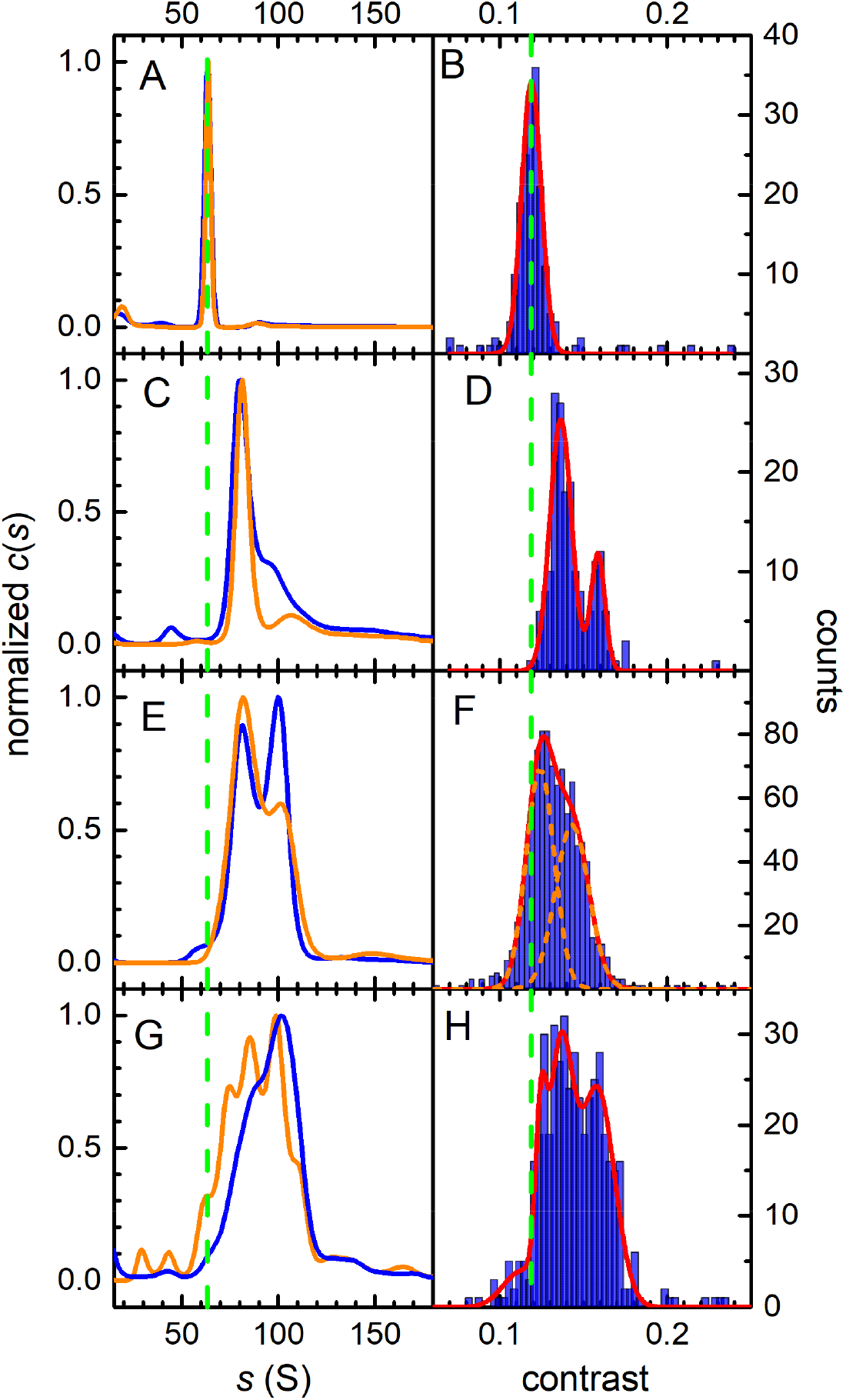
Comparison of the SV-AUC and MP data. (A, B) AAV8 empty capsids, (C, D) ssAAV8_4400_, (E, F) scAAV5-hSyn-GFP, (G, H) scAAV5-CMV-GFP. The left column shows SV-AUC *c*(*s*) distributions obtained from the 260 nm absorbance (blue) or Rayleigh interference (orange) signals, respectively. The right column shows MP contrast distribution histograms (blue) and their best fits with Gaussian distributions (red). Vertical dashed green lines show the position expected for empty capsid species in the *c*(*s*) and MP analysis, respectively. For the unresolved scAAV5-hSyn-GFP MP histogram in panel F, the two Gaussian fit components are shown as orange dashed lines.

### MP and SV-AUC analysis of partially purified rAAV preparations

To assess the practical performance of MP in the analysis of more complex samples, we compared the results of the SV-AUC and MP analysis of partially purified rAAV preparations. The SV-AUC data obtained for the empty AAV8 capsids reveal a single narrow *c*(*s*) distribution peak at the 64 S position, with a small amount of larger particles with *s* values in the 80 S to 100 S range (Fig. 5A). The MP histogram of this sample similarly shows a single narrow peak at the 1.19 contrast value with a small amount of particles detected in the higher contrast range (Fig. 5B). The *c*(*s*) distribution of the same rAAV8 stereotype with the encapsidated 4.4 kb ssDNA shows two main species—a dominant fraction at the 80 S position, and a smaller population with an approximate sedimentation coefficient value of 106 S (Fig. 5C). The MP distribution (Fig. 5D) correspondingly contains two well defined peaks with average contrast values of 1.36 and 1.58, and a relative abundance matching that shown by the *c*(*s*). Using the previously obtained contrast-to-nt calibration, the encapsidated genome sizes can be estimated at 2.1 ±0.2 kb and 4.3 ±0.2 kb for the left and right MP distribution peak, respectively. This suggests that the more abundant MP distribution peak indicates the presence of a large amount of vectors containing DNA impurities in this preparation, and the lower population peak represents the desired vector product.

SV-AUC of the scAAV5-hSyn-GFP vector containing the self-complementary DNA yields a *c*(*s*) distribution with two partially resolved peaks (Fig. 5E). The ratios of the interference and 260 nm absorbance signals indicate that the less abundant, higher *s*-value population contains a genome of a larger size. The MP histogram of the same sample does not completely resolve the peaks observed in the *c*(*s*) distribution (Fig. 5F). However, the increased peak width, and the well-defined shoulder of the distribution clearly indicate two underlying populations. Consequently, two Gaussian peaks were used to fit the MP distribution, and the sum of the two best-fit peaks describes the overall shape of the histogram very well. Using the previously established linear relationship between MP contrast and the DNA length, the genome size for the two peaks was estimated as 0.7 ±0.2 kbp and 1.8 ±0.1 kbp for the left and right peak, respectively. Based on these results, this scAAV5-hSyn-GFP preparation contains approximately 43% of the desired vector product and 57% of the encapsidated DNA impurities.

The final sample (scAAV5-CMV-GFP) encodes the same gene as the previously analyzed vector but contains a different promotor sequence. Nevertheless, the scAAV5-CMV-GFP SV-AUC data reveal a much more complex sample composition. The *c*(*s*) distribution indicates a small amount of empty capsids, represented by a shoulder at the approximately 60 S position and a wide distribution comprising of several species with *s* values spanning the 60 S to 120 S range (Fig. 5G). The ratio of the 260 nm absorbance to interference increases with the increasing sedimentation coefficient of the *c*(*s*) distribution species. This confirms that the wide SV-AUC distribution is reflecting the presence of several viral species encapsidating genomes of different sizes. The MP histogram revels a similarly complex picture (Fig. 5H). The conclusions that can be drawn from the analysis of the MP distribution are equivalent to those provided by the SV-AUC data. The low-contrast shoulder of the MP distribution extends over the 1.2 contrast region, where the empty capsids are observed. The overall width of the distribution is larger than that observed for other samples. Consequently, four Gaussian components are required to fit the MP histogram, and their combined envelope provides a good representation of the MP data. The genome sizes estimated from the position of the Gaussian peaks span the 1.1 kbp to 2.6 kbp range. Since the expected genome size of the scAAV5-CMV-GFP construct is 2.3 kbp, the MP analysis suggests that this vector preparation contains, besides the expected product, a small amount of empty capsids, and a large amount of virions encapsidating DNA impurities. Unlike the MP replicates of a highly purified sample (Fig. 2), the MP replicates of the scAAV5-CMV-GFP show more variability in the species populations detected in individual measurements (Fig. 6). This is most probably reflecting the fact that for a complicated, multi-species MP distribution, the particle count for each distribution species is relatively low. Each count represents an individual virion particle detected during the MP analysis. When multiple species are present, the total number of counts in the MP measurement is distributed over an increasing number of peaks, decreasing the accuracy of the measurement. Nevertheless, all MP replicate measurements show a similar pattern, in particular a comparable width of the distribution and the presence of multiple particle species. Their analysis leads to identical conclusions regarding the sample purity and the presence of the partially packaged capsid populations in the analyzed sample.

**Figure 6.**
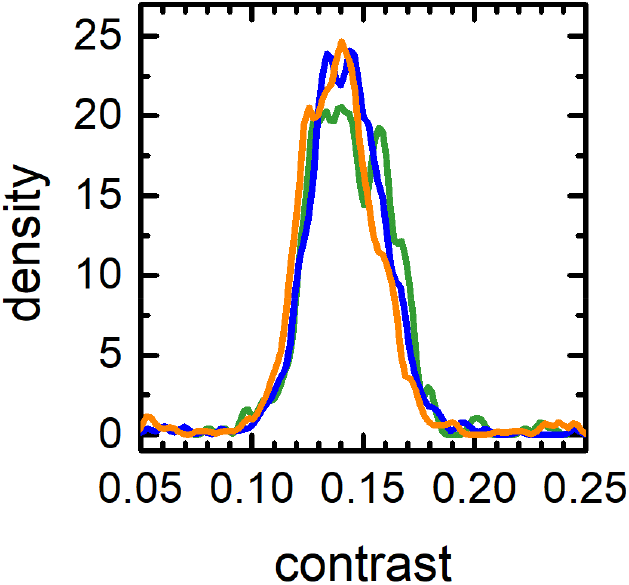
KDE plot of the triplicate MP measurements of the scAAV5-CMV-GFP sample shown in Figure 5H.

Taken together, the data indicate that MP provides extensive information on the heterogeneity of rAAV preparations. The resolution of the MP distributions is slightly lower than that obtained from the SV-AUC *c*(*s*) analysis, but MP is able to detect, identify, and characterize all species detectable by the *c*(*s*).

## Discussion

In this report, we present a comprehensive evaluation of the potential of MP as a technique to characterize rAAV vectors. Despite the fact that rAAVs have recently emerged as a potent new class of biopharmaceutical drugs, adequate methods to measure the purity and homogeneity of rAAV preparations are still lacking. MP has previously been shown to detect viral capsids, [33, 40], but this report is, to our knowledge, the first detailed study of MP as an rAAV analysis tool. Importantly, the MP results obtained here were validated by SV-AUC, a sensitive, high-resolution technique, able to quantify rAAV impurities not always detectable by other analytical methods. Our results show that MP contrast distributions provide information comparable to the *c*(*s*) SV-AUC data, but the data can be obtained faster and with minimal sample consumption. In contrast to AUC, a typical MP measurement requires a sub-picomole amount of sample and can be completed and analyzed within 5 minutes. This equals to approximately 1/600 of the sample and 1/40 of the time required by AUC. MP experiments are easy to execute—in our facility, using the commercial MP system—new users are typically able to independently collect and analyze MP data after approximately 30 minutes of training. Importantly, MP sample analysis is straightforward and easy to standardize. This guarantees that results obtained in separate experiments are comparable and not affected by arbitrary adjustments introduced by the operator.

The light scattering derived MP signal is generated by every particle in solution that has a molecular mass within the MP detection range. This lack of signal source discrimination can be problematic in some MP applications. It precludes the MP analysis of samples with a large amount of impurities, particularly when the molecular mass of the impurities is similar to that of the species of interest. However, this MP attribute is advantageous for sample quality assessment applications. For the same reason, best MP results are obtained from the analysis of fresh vector preparations. We have observed that repeated freeze-thaw cycles substantially increase the amount of vector debris and degradation products in the sample solution. Light scattering generated by those impurities can quickly dominate the MP signal. The MP data analysis parameters used in this work suppress the detection of impurities in the sub-MDa range (see Methods section). However, the raw MP data contain complete information on all detected particles. If the small molecular mass impurities are of interest, filter parameters can be adjusted and MP data reanalyzed to include all molecules detected in the full MP sensitivity range. Recently developed upgrades of the MP controller software may further improve the AAV data collection. Since viruses and other large molecules diffuse slowly, the frequency of landing events detected by the MP instrument for those molecules is relatively low. The upgraded controller software can expand the size of the MP field of view (FOV) to increase the number of rAAV molecules measured in a single data acquisition session. At the same time, larger FOV will somewhat decrease the sensitivity of the MP instrument to small molecules. All data collected for this manuscript were acquired before those upgrades became available. Since MP is a relatively new technology, future technical advancements may further improve MP capabilities.

The most straightforward MP application for rAAV analysis is the assessment of rAAV sample heterogeneity. Figure 5 illustrates the information content of MP contrast distributions for increasingly complex rAAV samples. Clearly, when relatively pure samples are analyzed, MP can quickly confirm their homogeneity (Fig. 5B) or identify the nature and the fractional content of the molecular components (Fig. 5D, 5F). Obtaining precise component information for a more complex, heterogenous sample is difficult (Fig. 5H), but, nevertheless, the MP data matches the information obtained from other methods (Fig. 5G).

Besides the assessment of sample homogeneity, the relative numerical ratios of sample populations can also be easily obtained by Gaussian fitting of the MP contrast distributions. For samples with empty and full capsid populations, this allows for quick and accurate packing efficiency determination (Fig. 3). These calculations are unambiguous for samples with limited heterogeneity (Fig. 3A, 5D) but become increasingly more difficult for data with partially resolved species (Fig. 5F) and with multiple species populations (Fig. 5H). From our data, we estimate that MP can typically resolve rAAV species with genome size differences of approximately 0.7 kb. The resolution will also be affected by the relative size of the MP distribution peaks—populations with a relatively low particle count can be detected by MP, but their precise numerical analysis will be limited. Finally, the genome size can be estimated from the species Gaussian peak position. However, this requires obtaining the contrast-to-nt calibration information from separate measurements of standard samples.

The area under the MP distribution plots is equal to the total number of detected virions. This number will correspond to the concentration of virions in solution, and could potentially be used for titer estimations by comparing the sample particle count with that of an appropriate concentration standard. There are, however, several factors that can influence the accuracy of this measurement. Some of the landing events can be lost in the background noise if the mass photometry images are of poor quality. The mass photometry optimum concentration range is relatively narrow, and the concentration of the analyzed sample will typically have to be adjusted to obtain best results. The strong negative charge of viral DNA genomes could also potentially affect the virion’s propensity to land on the surface of the glass coverslip, especially in low ionic-strength buffers. All those factors would have to be carefully explored, but it is feasible that a reliable mass photometry protocol for vector titer estimation could potentially be developed.

This study shows that MP can measure heterogeneity, relative species content, packing efficiency, and other attributes of AAV preparations accurately and reproducibly. MP measurements are fast, sensitive, inexpensive, and typically require no sample manipulation other than the concentration adjustment. Those attributes make MP an attractive tool for the biophysical characterization of viral vectors in industry and academic applications.

## Acknowledgements

This work was supported by the intramural program of the NHLBI, NIH.

---

**Supplemental Figure 1.**
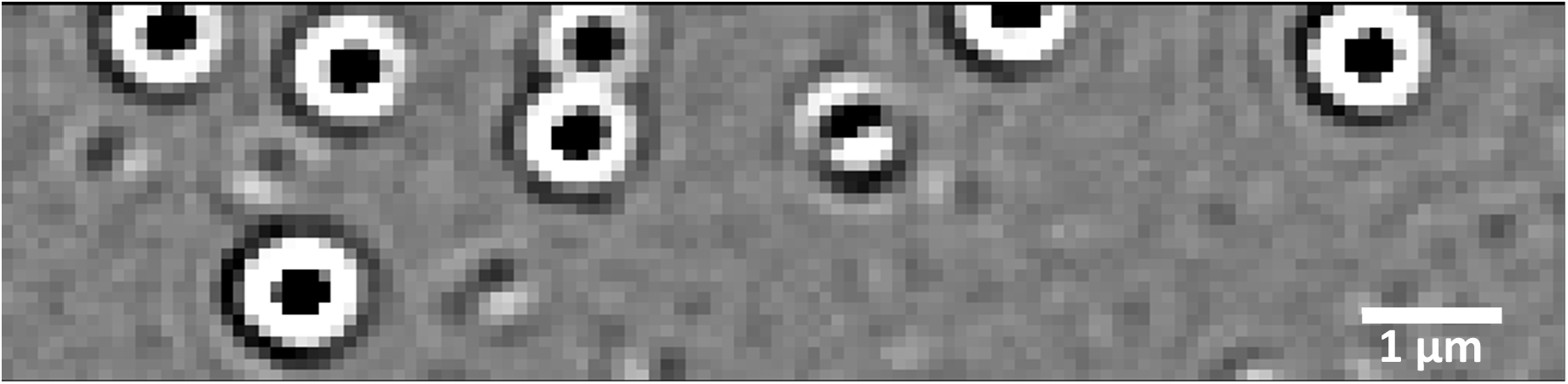
MP image frame of a rAAV sample

## Notes

### Competing Interest Statement

The authors have declared no competing interest.

